# A locally administered single cycle influenza vaccine expressing a non-fusogenic stabilised haemagglutinin stimulates strong T-cell and neutralising antibody immunity

**DOI:** 10.1101/2024.03.01.582898

**Authors:** Holly L. Sadler, Pramila Rijal, Tiong Kit Tan, Alain R. M. Townsend

## Abstract

Current influenza vaccination approaches protect against specific viral strains, but do not consistently induce broad and long-lasting protection to the diversity of circulating influenza viruses. Single cycle viruses delivered to the respiratory tract may offer a promising solution as they safely express a diverse array of viral antigens by undergoing just one round of cell infection in their host and stimulate broadly protective resident memory T-cell responses in the lung. We have previously developed a vaccine candidate called S-FLU that is limited to a single cycle of infection by inactivation of the hemagglutinin signal sequence and induces a broadly cross-reactive T-cell response and antibodies to neuraminidase, but fails to induce neutralising antibodies to hemagglutinin after intranasal administration.

This study describes the development of CLEARFLU, a derivative of S-FLU that is designed to add a neutralising antibody response to hemagglutinin. In contrast to S-FLU, which does not express a hemagglutinin molecule at the infected cell surface, CLEARFLU viruses express a stabilised non-fusogenic hemagglutinin. They are equally limited to a single cycle of infection, but induce a neutralising antibody response to the expressed hemagglutinin in addition to the cytotoxic T lymphocyte (CTL) responses to internal proteins and antibodies to neuraminidase induced by S-FLU. This represents a notable advantage as CLEARFLU viruses may provide sterile immunity against strain-matched challenge as well as non-sterile protection against a broad range of influenza viruses.

**Importance:** Influenza is a serious public health concern, causing seasonal epidemics as well as pandemics in people. Influenza can also cause severe agricultural losses due to its circulation in farmed poultry and swine. A major challenge in the control of influenza is the diversity of circulating viruses. Developing vaccines which stimulate immunity to a wide array of influenza viruses is therefore important for protecting human and animal populations from disease and death. In this study, we describe an approach for developing influenza vaccines which trigger immune mechanisms shown to induce broad protection against a diversity of viruses, while also conserving the strong protection against specific strains observed in existing vaccines.

## Introduction

Viruses that undergo a single replication cycle offer a promising platform for generating broadly protective immune responses to influenza (1–3). They are designed to lack a functional copy of one or more of the genes required for influenza replication and can only be propagated when the related gene product is supplied in *trans*. While single cycle viruses can be propagated to high titre in permissive cell culture, they are able to undergo only a single cycle of cell infection in their natural hosts.

Single cycle influenza viruses (SCIV) with a functioning polymerase undergo genome replication and expression in their host, leading to amplification of viral epitopes inside infected cells and cell surface expression of encoded viral coat proteins. As SCIV typically contain all but one intact viral segment, responses can be targeted to a wide array of RNA, peptide and protein components. SCIV are therefore capable of generating strong antibody and T-cell responses, including broadly protective cytotoxic T-cell responses to peptides from highly conserved internal core proteins (4). SCIV hold an advantage over conventional live-attenuated vaccines in that they infect very few cells and so are unlikely to cause pathology or mutate or reassort into virulent forms. SCIV can be safely administered by aerosol to immunise the lung, which is associated with local immunity through tissue-resident T-cells (5–7) and mucosal IgA (8).

Multiplasmid transfection is commonly used to produce influenza viruses *de novo* in cell culture (9–11). To generate single cycle viruses, one or more producer plasmids can be mutated or omitted. SCIV vaccine candidates with deficiencies in various genes have been developed, including in matrix 2 (3), neuraminidase, the polymerase genes and the hemagglutinin. Inactivating the hemagglutinin is particularly advantageous as it prevents functional hemagglutinin reassortment. Hemagglutinin activity is also easy to compensate for *in trans* by growing viruses in cells which express functional coating hemagglutinin on their surface. Altering the coat protein in the producer cell lines (pseudotyping) can allow single cycle viruses with the same core genes but varying coating hemagglutinins to be produced quickly from a single seed virus. This has the advantage that the coating hemagglutinin can be selected to avoid prior immunity and enhance the cell mediated immune response (12).

We have previously described the single cycle virus S-FLU, which is produced by inactivation of the hemagglutinin signal sequence, generating a viral RNA called S-HA (signal minus hemagglutinin) (13). Like for most other SCIVs lacking hemagglutinin activity, no hemagglutinin epitopes are presented on the surface of cells infected with S-FLU. In other designs, the majority of the hemagglutinin coding sequence can be replaced by other genes; for example fluorescent proteins to follow cell infection (14), or NY-ESO-1 to induce tumour immunity (12). S-Flu viruses have been helpful laboratory tools for evaluating the susceptibility of different coat proteins to chemical inhibitors and antibodies. Recently, S-FLU viruses coated in avian H7 hemagglutinins (15) and the glycoprotein from Ebola viruses (16,17) have been evaluated as safe and accurate pseudotypes for assaying inhibition of cell entry. This enables the characteristics of coat proteins from highly virulent viruses to be investigated in biosafety containment 1/2 rather than in high containment facilities.

S-FLU viruses have shown promise as broadly protective vaccines in mice, ferrets and pigs. They can be administered into the lung intranasally (mice and ferrets) or by aerosol (pigs) without causing pathology. Heterologous protection is induced in the absence of a neutralising antibody response to hemagglutinin at the low doses administered to the lung. Protection is associated with strong T-cell responses, including CD8+ T-cell responses in the lung (7), and a strain-specific antibody response to neuraminidase which reduces challenge virus titres in the respiratory tract in mice (13,15) and ferrets (4,18) and may (19) or may not (18) reduce challenge virus titres in pigs. No sterile immunity is evoked in any model species following administration of S-FLU to the lung. While S-FLU viruses protect against severe disease, they therefore are not able to prevent infection, even if they are perfectly antigenically matched to the challenge strain.

In this study, we explore whether expressing hemagglutinin epitopes from a non-functional hemagglutinin molecule instead of S-HA or eGFP in the S-FLU expression cassette drives their cell surface expression and stimulates neutralising antibody responses without negatively impacting on the existing immunogenicity and single cycle nature of S-FLU viruses.

A multitude of single or double nucleotide mutations which significantly ablate or abolish hemagglutinin function have been described previously. While alone these mutations generate single cycle viruses with a high rate of reversion to infectivity, using them in combination could allow for the generation of a full length, rationally designed, multi-mutated hemagglutinin molecule which is irreversibly non-functional but antigenically preserved.

Here, we describe the iterative design of a non-functional multi-mutated hemagglutinin immunogen named CLEARFLU. We show that single cycle influenza viruses expressing CLEARFLU hemagglutinin generate neutralising antibody responses as well as broadly reactive T-cell responses in mice, when administered at low dose to the lung.

## Results

### CLEARFLU design and expression

The design of CLEARFLU hemagglutinins was refined through three iterations using H1, H3 and H7 hemagglutinins (Table 1). Each CLEARFLU design included seven independent sets of mutations known to block or heavily ablate hemagglutinin fusion activity, acting through up to five different mechanisms: resistance to proteolytic cleavage (20,21), conformation locking through disulphide bonding (22,23), receptor inactivation (24,25), inhibition of the fusion peptide (26) and B-loop inactivation (27).

**Table 1.**
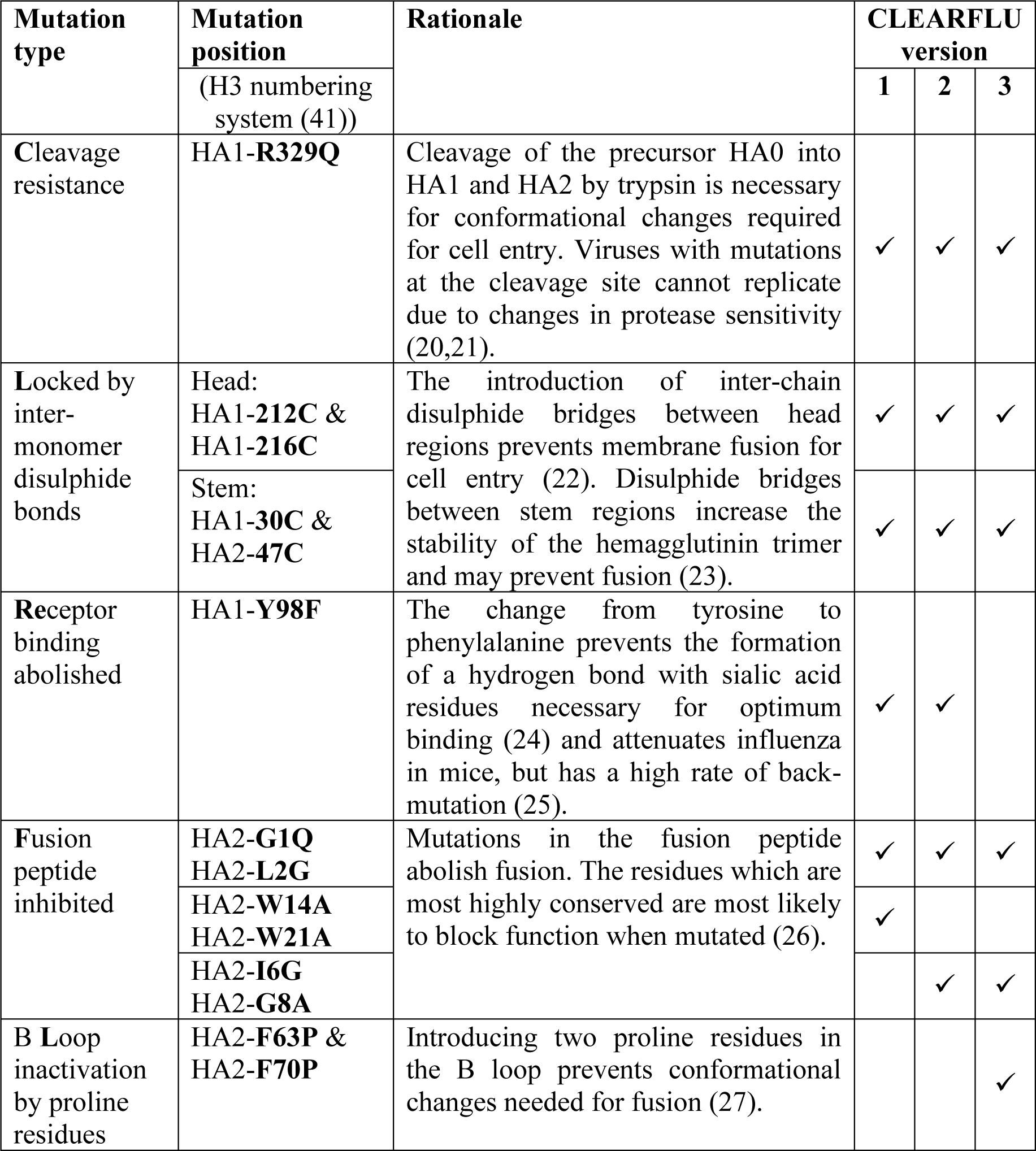
Mutations introduced to inactivate wild-type hemagglutinin genes in each version of the CLEARFLU design.

In the first design, the selected mutations disrupted binding by some antibodies (T1-3B, T3-5D, T2-6D (28) and MEDI8852 (29)) to the stem of H1 and H3 hemagglutinins expressed in cell lines (Figure 1). This binding was restored for CLEARFLU version 2 by reverting mutations W14A and W21A to wild-type and replacing these with I6G and G8A, which also inactivate the fusion peptide but are closer to the N terminus and do not interfere with the binding site for these antibodies. Antibody binding to the hemagglutinin globular head did not appear to be affected by the introduction of any candidate mutations.

**Figure 1.**
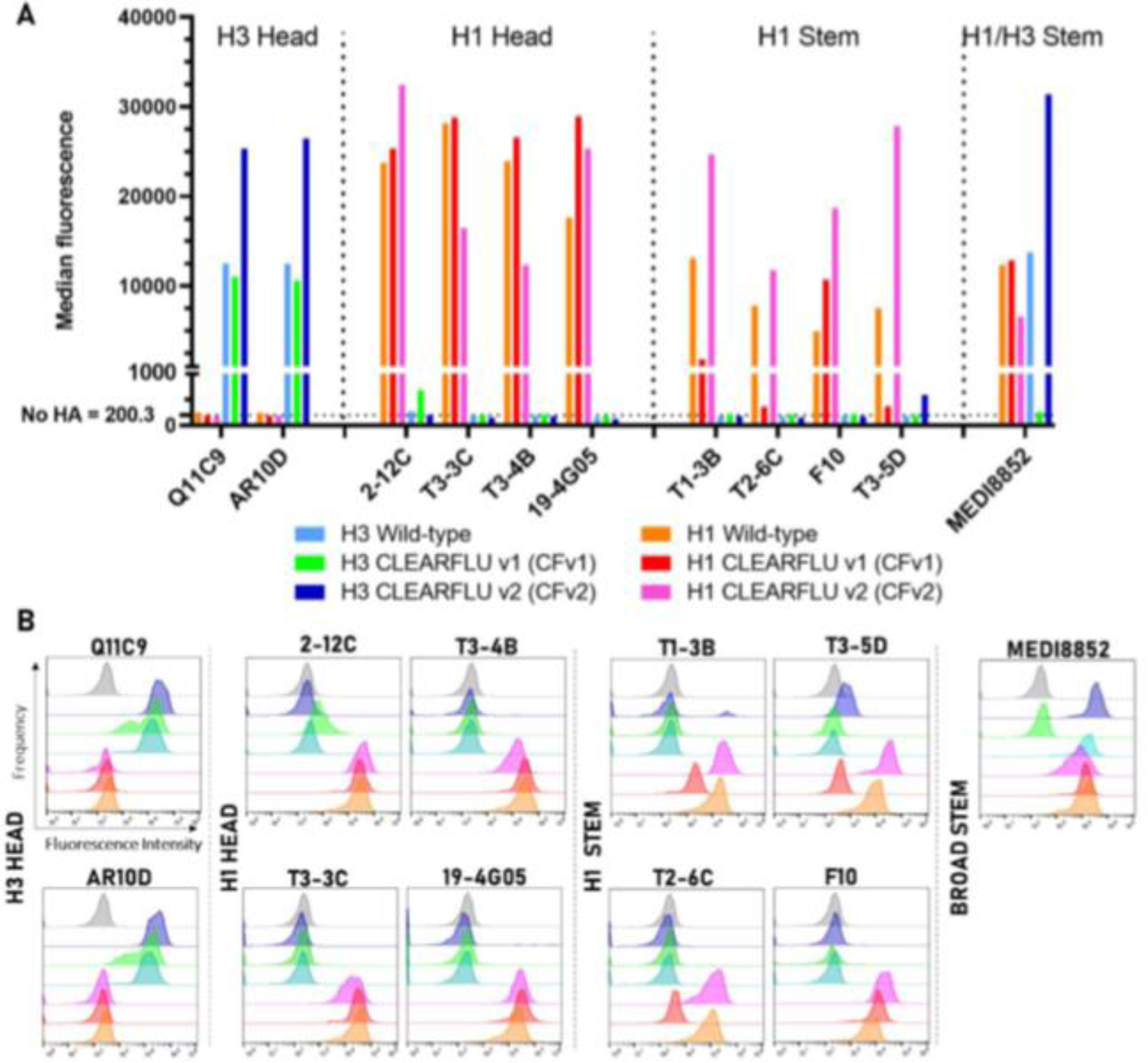
Antibody binding panel to CLEARFLU versions 1 & 2 expressed in stably transduced MDCK-SIAT1 cells. MDCK-SIAT1 cells were transduced to express H1 A/England/195/2009 and H3 A/Hong Kong/5738/2014 CLEARFLU and wildtype hemagglutinins and sorted for high expression with antibodies T3-3C (28) (H1) or AM3C (H3) (produced in-house, manuscript in preparation). Hemagglutinins were bound with head or stem-targeted human primary antibodies, and binding was detected with a FIT-C labelled goat anti-human IgG secondary antibody. A. Bar graph showing median fluorescence. The average median fluorescence for three populations of unstained cells which were not transduced to express hemagglutinin is shown as the no-HA control. B. Histogram plots showing the relative frequency of cells detected with a given fluorescence intensity. Grey plots show unstained cells which were not transduced to express hemagglutinin.

To verify that the mutations in the CLEARFLU version 2 design did indeed inactivate hemagglutinin activity, we tested the replication of S-FLU viruses in cell lines expressing H7 A/Hong Kong/125/2017 hemagglutinins each with a single inactivating mutation (see Figure 2). As S-FLU viruses express eGFP instead of hemagglutinin, green comma-shaped plaques of infected cells are formed only in cell lines expressing functional hemagglutinin. We found no evidence that the Y98F mutation previously described to inactivate the sialic acid binding site reduced the growth of S-FLU in H7 transduced MDCK-SIAT1 cells. This supports previous observations by others that Y98F leads to reduced receptor binding, but only limits infection in cells with low sialic acid expression (30,25). As MDCK-SIAT1 cells over express α-2,6-sialic acids, this effect was not seen here (31). The G8A mutation appeared to significantly reduce the size of viral plaques, and we did not observe peaks of fluorescence in middle dilutions to suggest efficient expansion of single viral clones. This suggests that the intermediate fluorescence recorded for G8A is due to many viruses replicating poorly and that G8A reduces but does not abolish fusion activity in this context. All the other mutations appeared to completely abolish hemagglutinin-dependent replication.

**Figure 2.**
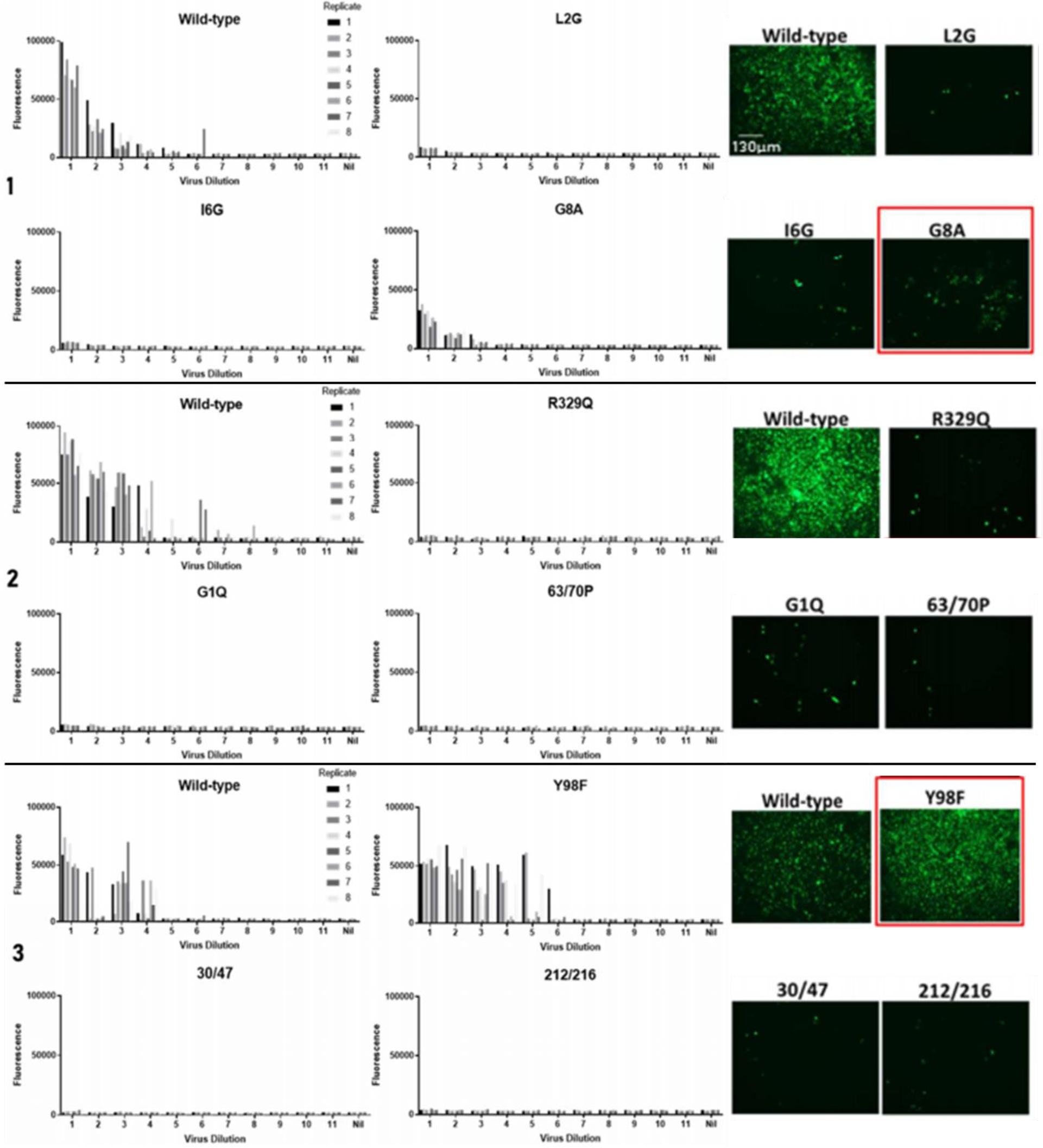
Replication of S-FLU virus in cell lines expressing hemagglutinin with a single CLEARFLU candidate mutation. MDCK-SIAT1 cells were transduced to express the selected mutant H7 A/Hong Kong/125/2017 hemagglutinin and sorted for high expression with H7 antibody 4A14 (Huang 2019). Cells were infected with an eGFP-expressing S-FLU virus pseudotyped with functional hemagglutinin in half-log dilutions, starting at 1000 TCID50 per well. Eight replicates were used at each dilution. Bars show the fluorescence of each well in a 96-well plate after 48 hours. Fluorescence microscopy images show cells in dilution 1 expressing eGFP after S-FLU infection, with clusters of infected cells indicating virus replication. The experiment was split into three sets, each of which was repeated with the same results. Virus used for infection: H7N1 S-FLU ([S-eGFP/N1(A/Puerto Rico/8/1934)] coated in H7(A/Netherlands/219/2003))

To further refine the design of CLEARFLU, we removed the Y98F mutation to avoid disrupting untested neutralising epitopes around the receptor binding site. To maintain the number of inactivating mutations, we introduced two proline residues at positions 63 and 70 in HA2, which together prevent conformational changes in the B loop required for fusion (27) and are not permissive of S-Flu replication (Figure 2). CLEARFLU version 3 thus contains an intact receptor binding site (Y98), seven fully inactivating mutation sets (R329Q, G1Q, L2G, I6G, 30/47C, 212/216C, 63/70P) and the partially attenuating mutation G8A. An antibody panel specific for H7 haemagglutinin (32) confirmed that this combination of mutations did not disrupt protein folding (Figure 3).

**Figure 3.**
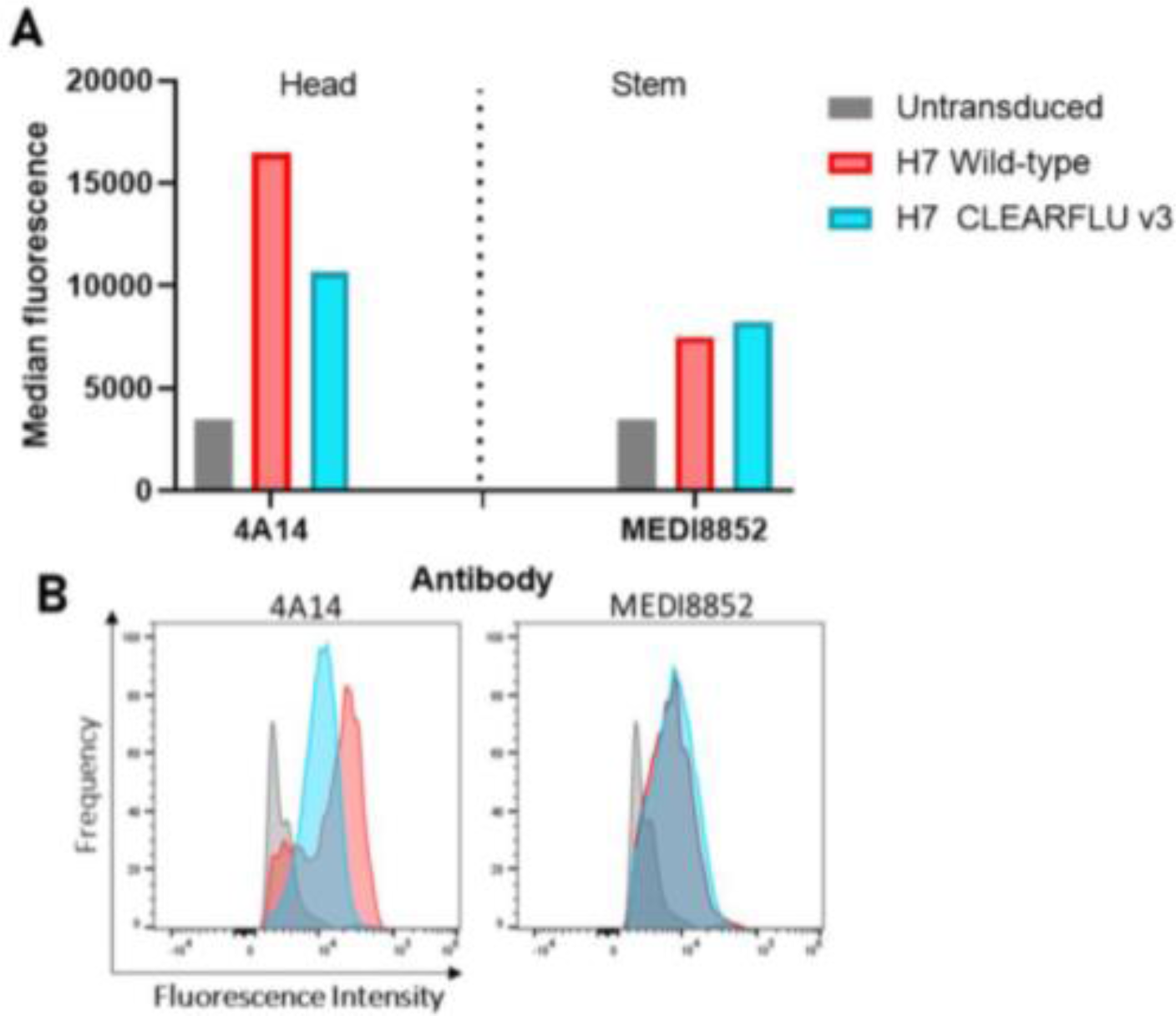
Antibody binding to CLEARFLU version 3 expressed in stably transduced MDCK-SIAT1 cells. MDCK-SIAT1 cells were stably transduced to express H7 A/Hong Kong/125/2017 CLEARFLU version 3 hemagglutinin and sorted for high expression with H7 antibody 4A14. Hemagglutinins were bound with head or stem-targeted human primary antibodies, and binding was detected with an Alexa Fluor 647 labelled secondary antibody. A. Bar graph showing median fluorescence. B. Histogram plots showing the relative frequency of cells detected with a given fluorescence intensity.

### Genomic stability of CLEARFLU viruses

Viruses expressing H7 A/Hong Kong/125/2017 CLEARFLU version 3 in the hemagglutinin expression cassette were generated by multiplasmid transfection of HEK 293T cells (9). These viruses were expanded in MDCK-SIAT1 cells stably expressing H1 A/Puerto Rico/8/1934 functional hemagglutinin.

CLEARFLU viruses grew to titres >10E6 TCID50/ml / >10E7CID50/ml (see methods), but while CLEARFLU was strongly displayed on the surface of some infected cells, many viral plaques did not stain with H7-specific antibody 4A14 (32), which targets the receptor biding site (Figure 4). This suggested that CLEARFLU HA expression had been lost or that the receptor binding site had been disrupted. The loss of full length CLEARFLU during expansion may not be surprising given that CLEARFLU expression is not expected to confer any replicative advantage in these conditions.

**Figure 4.**
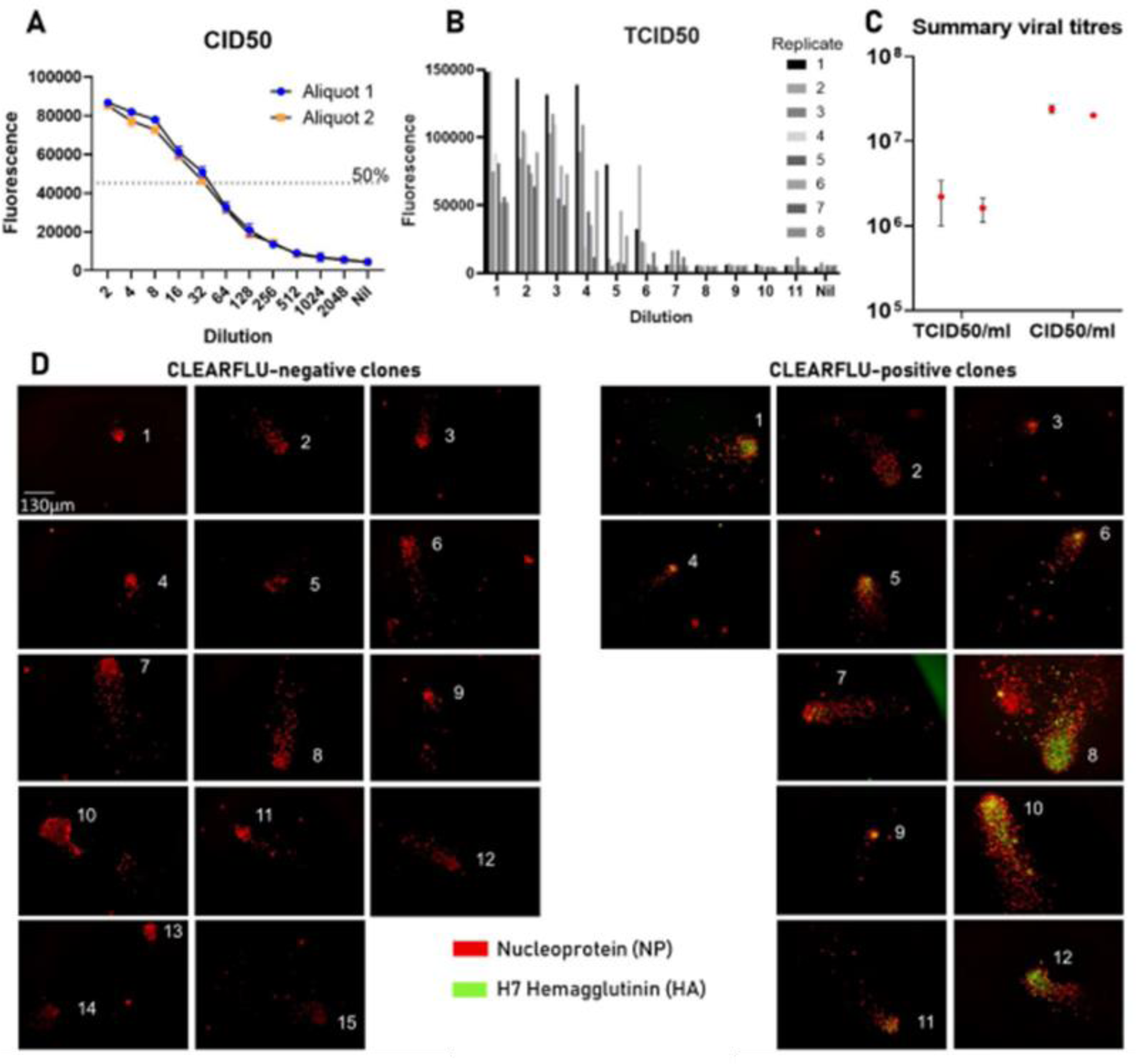
Titration and hemagglutinin expression of CLEARFLU version 3 viruses. A, B & C. Two aliquots of CLEARFLU version 3 virus reference stock were titrated by CID50 and TCID50 using antibody 2-8C (Powell et al., 2012) specific to nucleoprotein and an Alexa Fluor 647 labelled secondary antibody. Error bars show the standard deviation around the mean from duplicates. Plot B is representative of two titrations for each aliquot, the results of which are presented in C. D. Fluorescence microscopy images show the expansion of CLEARFLU version 3 virus clones in MDCK-SIAT1 cells transduced to express PR8 hemagglutinin. Cells were infected as per TCID50 protocol but were stained after 24 hours with primary antibodies 2-8C (biotin-conjugated) and 4A14 and then with secondary layers Alexa Fluor 647 streptavidin and goat anti-human Alexa Fluor 488. Clones were defined as clusters of nucleoprotein-stained cells, often with comet-like trails.

Previous work in our laboratory has shown that S-FLU viruses grow relatively poorly in MDCK-SIAT1 cells expressing Ebola glycoprotein (GP) (16). We therefore trialled expanding CLEARFLU viruses in cells expressing GP to determine whether this would select for expression of the intact hemagglutinin receptor binding site in CLEARFLU version 3 by providing an efficient sialic acid binding function.

CLEARFLU viruses indeed grew to higher titres and formed denser plaques in MDCK-GP cells than S-FLU viruses, and plaques produced by CLEARFLU viruses which did not stain with hemagglutinin antibody were similar in morphology to small diffuse plaques produced by S-FLU viruses (Figure 5). We tested this apparent selective pressure and genomic stability under these conditions by passaging the virus at low multiplicity of infection (MOI) six times in MDCK-GP cells, representing a 4x10^18^ expansion by round six. After six rounds, CLEARFLU viruses still grew to high titre and we were able to detect CLEARFLU expression, however the proportion of clones which stained with 4A14 decreased from over 90% to around 10% (Figure 6). Remarkably, sequencing revealed that no mutations were identified in the coding sequences of any core viral genes in the round six culture.

**Figure 5.**
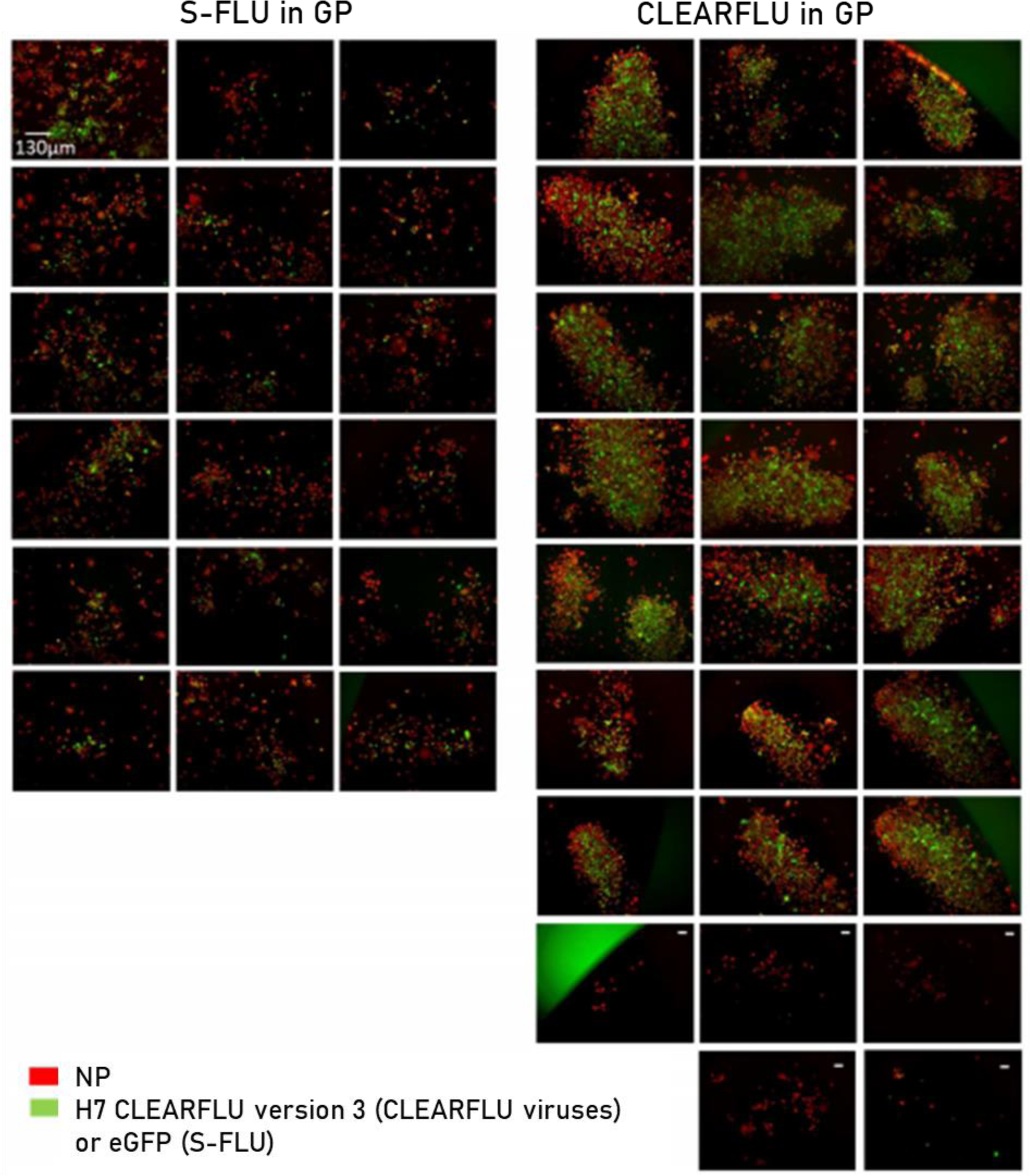
Plaque morphology of CLEARFLU version 3 viruses grown in cells expressing Ebola glycoprotein (GP). Cells were infected with S-FLU or CLEARFLU version 3 virus as per TCID50 protocol and stained after 48 hours with anti-nucleoprotein mouse antibody AA5H and an Alexa Fluor 647 secondary antibody (red). Cells infected with CLEARFLU version 3 virus were also stained with H7 hemagglutinin human antibody 4A14 and an Alexa Fluor 488 secondary antibody (green). Fluorescence microscopy images were taken of plaques indicative of clonal expansion. CLEARFLU virus plaques which did not stain with 4A14 are indicated with a white dash. Viruses: H7N1 S-FLU ([S-eGFP/N1(A/Puerto Rico/8/1934)] coated in H7 A/Hong Kong/125/2014) & H7 A/Hong Kong 125/2014 CLEARFLU version 3 reference stock.

**Figure 6.**
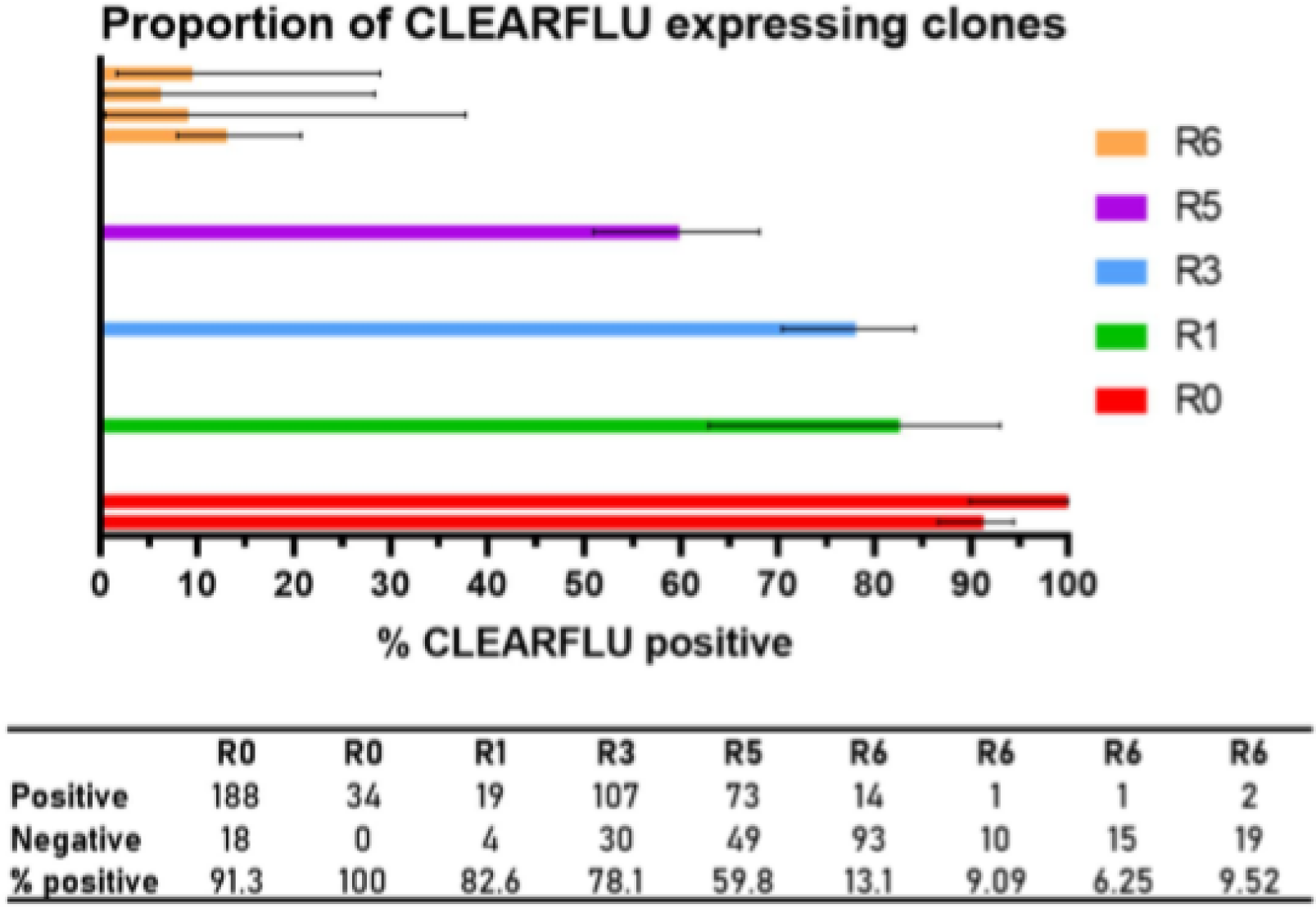
Proportion of clones expressing CLEARFLU version 3 hemagglutinin after passage at low MOI in MDCK-SIAT1 cells expressing Ebola glycoprotein. CLEARFLU version 3 reference stock was used to infect a large flask of MDCK-SIAT1 cells expressing Ebola GP at a multiplicity of infection of around 0.01 by TCID50, and then passaged at a 1 in 1000 dilution every 48 hours six times. Aliquots from each round were used to infect MDCK-SIAT1 cells expressing GP in limiting dilution, and viral clones were identified by nucleoprotein staining with AA5H (Alexa Fluor 647 secondary). Clones were scored as positive for CLEARFLU expression if they also stained with 4A14 (Alexa Fluor 488 secondary). Each bar shows an independent count. Error bars show 95% confidence intervals calculated using the Wilson/Brown method for proportions.

In isolating viral clones from the round six culture, we were able to identify several CLEARFLU viruses which replicated efficiently to form dense plaques despite not staining with 4A14. Four of these clones were sequenced and all were found to carry the T148A mutation (N2 numbering; T132A in N1 numbering) in their neuraminidase. The T148A substitution has been previously reported to disrupt a conserved site for N-linked glycosylation in the B1L23 loop [HS**NGT**VKDR] close to the active site of the enzyme (33).

To confirm that this substitution was responsible for the enhanced growth in these conditions, we introduced the T148A mutation alone into S-FLU. We showed that this mutation enables the neuraminidase to agglutinate red blood cells and enhances cell entry in the GP transduced MDCK-SIAT1 cells in an oseltamivir and neuraminidase antibody sensitive manner (see S.12 and S13 in the supplementary material). This suggested that T148A enhanced cell entry by enhancing the binding to sialic acid, possibly reducing the selection for expression of CLEARFLU. Sequencing of the CLEARFLU HA segment of two T148A carrying clones revealed a large truncation of 1620 nucleotides, leaving a 438 nucleotide fragment containing intact packaging sequences but only 63 nucleotides of CLEARFLU coding sequence (Figure 7). In another clone which grew well, stained with 4A14 and did not contain the T148A mutation, the CLEARFLU segment remained full-length and contained no mutations.

**Figure 7.**
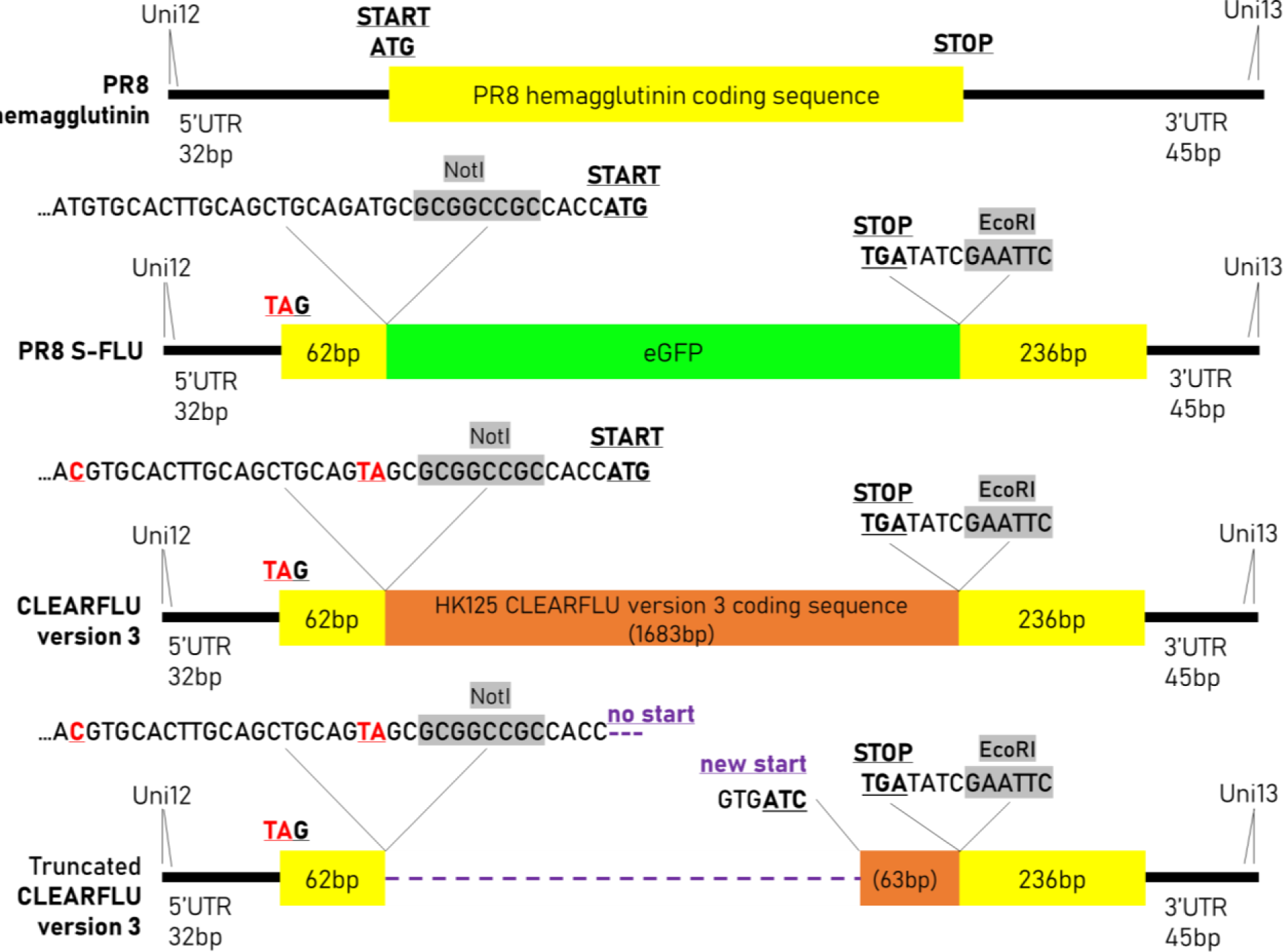
Design of the CLEARFLU version 3 viral expression cassette and truncation after repeated passage in MDCK-SIAT1 cells expressing Ebola glycoprotein. The CLEARFLU expression cassette contains intact packaging signals from the S-FLU expression cassette, which in this design differs from that described in Powell et al. 2012 by the alteration of two additional interfering ATG codons upstream of the NotI site. After six rounds of passage at low multiplicity of infection in MDCK-SIAT1 cells expressing Ebola glycoprotein, two viral clones expressed a truncated CLEARFLU construct containing only 63 base pairs of coding sequence flanked by intact PR8 untranslated regions (UTRs).

These results showed that expressing the CLEARFLU HA can provide a growth advantage (presumably by enhancing sialic acid binding) to viruses propagated in cell lines transduced with Ebola GP to provide a fusion function. However, substitution of NA T148A can also provide enhanced sialic acid binding, and viruses with this mutation compete with those expressing CLEARFLU HA.

### Murine immune response to CLEARFLU viruses

As the immune response to S-FLU viruses is known to protect mice, ferrets and pigs against heterologous challenge, we compared the immune response to CLEARFLU and S-FLU viruses in mice. In our first experiment, we assessed the serum antibody response to viruses expressing seasonal H1(A/England/195/2009) and H3(A/Hong Kong/5738/2014) CLEARFLU version 2 hemagglutinins. In the second, we compared local and systemic, humoral and cellular immune responses to a virus expressing H7(A/Hong Kong/125/2017) CLEARFLU version 3 hemagglutinin to those generated by a strain-matched S-FLU.

Like S-FLU, CLEARFLU version 2 and CLEARFLU version 3 viruses can be safely administered to mice both intranasally and intraperitoneally. We found that CLEARFLU viruses offer an advantage over S-FLU viruses in that they produce a strong neutralising antibody response to hemagglutinin even when administered to the airways (Figures 8 & 9). This response was particularly strong for H1 and H7 hemagglutinins compared to H3. For CLEARFLU version 3, this neutralising response was strongest in the serum, but also detected in bronchoalveolar lavage (BAL) and, while subtype-specific, showed some within-subtype cross-neutralisation in the serum (Figure 9). The antibody response to the neuraminidase in the BAL and serum was similar to that generated by S-FLU (15).

**Figure 8.**
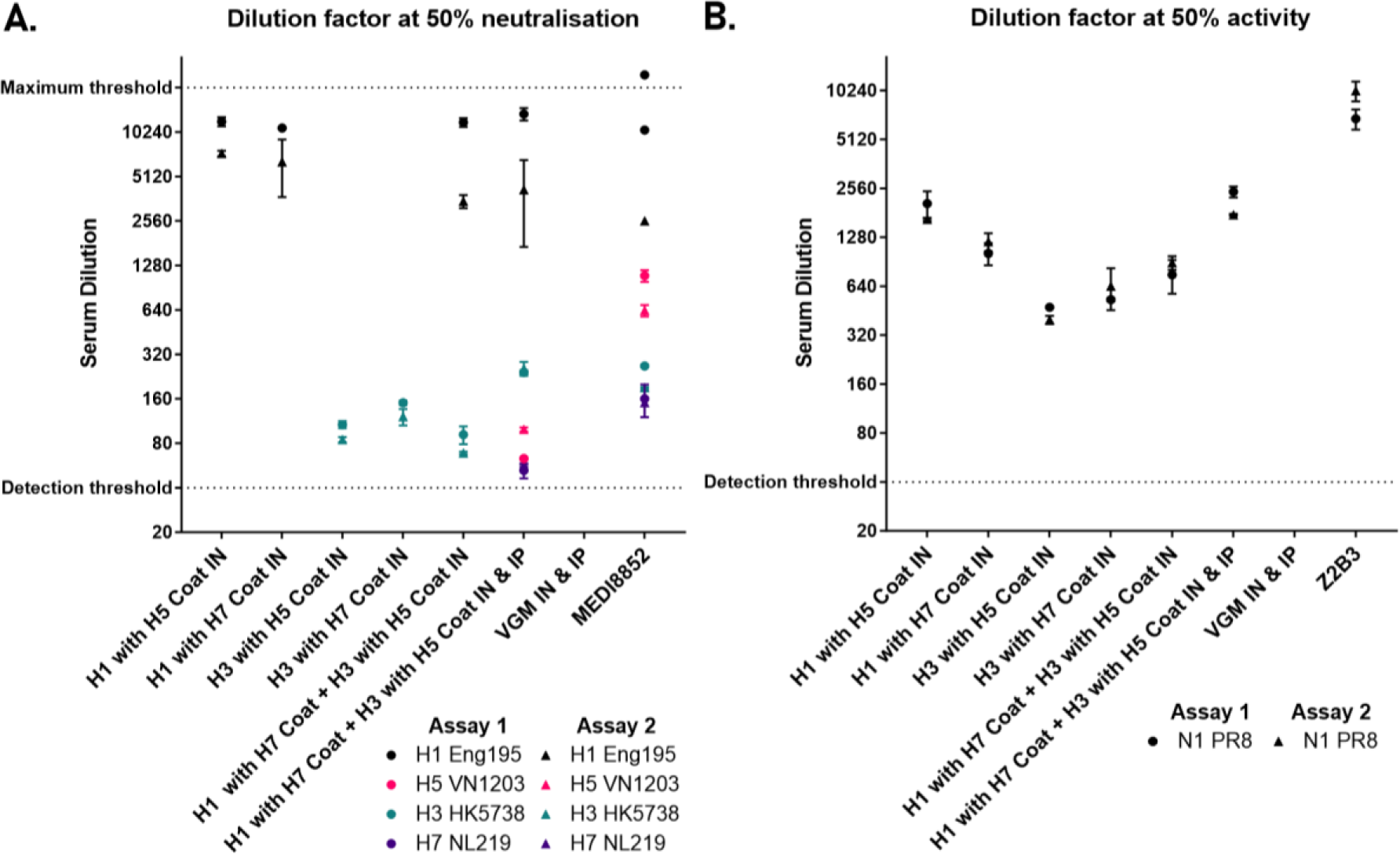
Murine serum antibody response following immunisation with CLEARFLU version 2. Serum antibody response in mice immunised with H1 or H3 CLEARFLU version 2 viruses pseudotyped with H5 or H7 coats. Mice were immunised with 2E6 CID50 virus intranasally (IN), or 1E6 CID50 intranasally and 1E7 CID50 intraperitoneally (IP) twice 28 days apart. Sera were harvested 14 days after the second dose for characterisation by microneutralisation (A) and enzyme linked lectin assay (B). Sera from 5 mice were pooled for each group, except the viral growth medium (VGM) only control group, for which 4 mice were used. Error bars show the standard deviation from within-assay duplicates, with results from two independent assays shown. A. Neutralising antibody response to H1, H3, H5 and H7 S-FLU by immune mouse sera or control antibody MEDI8852 B. Inhibition of N1 neuraminidase activity by mouse immune sera or control antibody Z2B3.

**Figure 9.**
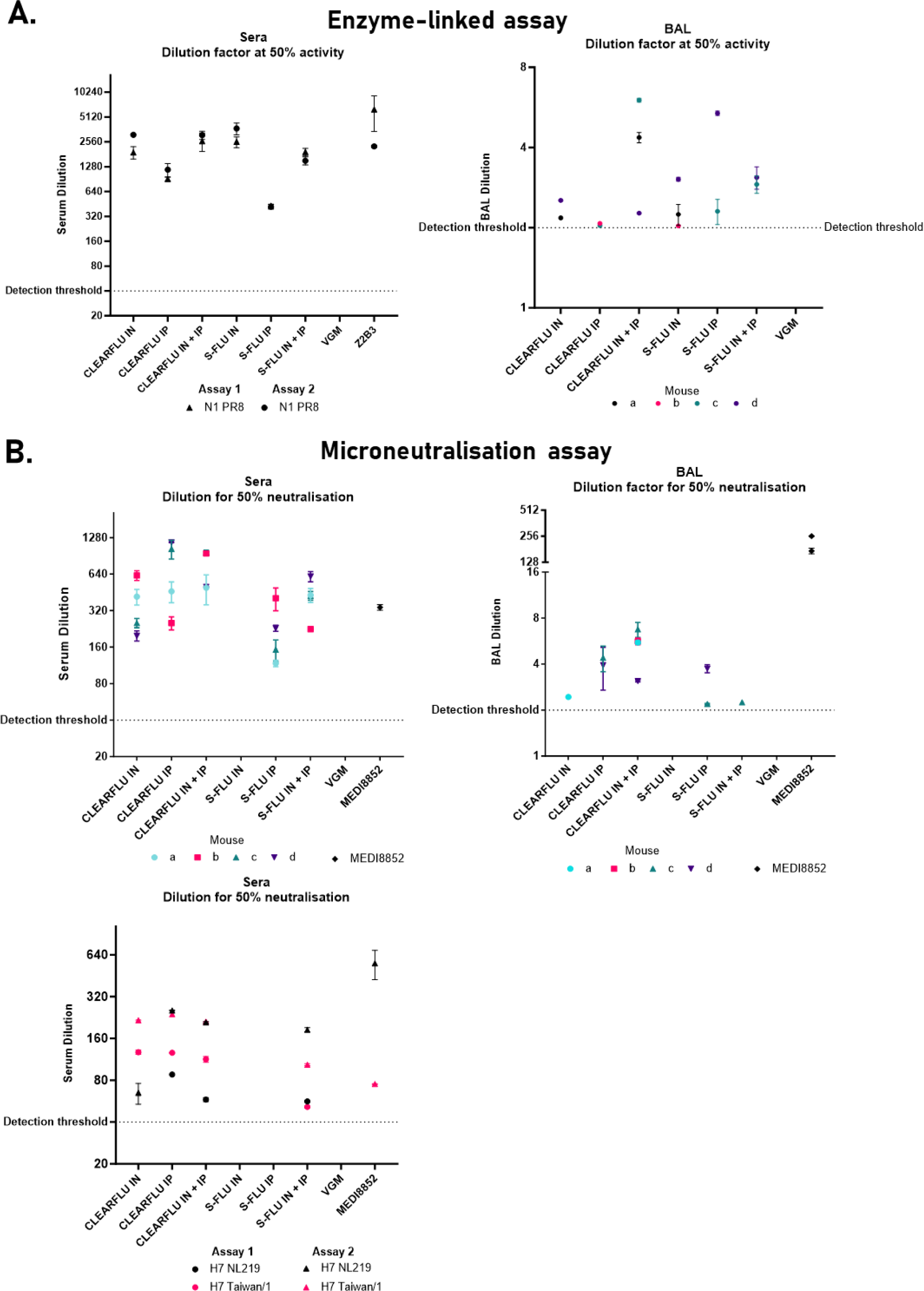
Murine serum and bronchoalveolar lavage (BAL) antibody response following immunisation with CLEARFLU version 3. Serum and BAL antibody response in mice immunised with H7 CLEARFLU version 3 pseudotyped with an H1 (PR8) coat or a strain-matched H7 S-FLU. Mice a-d in each group were immunised twice 14 days apart with 1E5 TCID50 virus intranasally (IN), 1E6 TCID50 intraperitoneally (IP) or 1E5 TCID50 IN and 1E6 TCID50 IP. Sera and BAL were harvested 68 days after the second dose for characterisation by enzyme-linked lectin assay (A) and microneutralisation (B). Error bars show the standard deviation from within-assay duplicates when both duplicates fell above the detection threshold. A. Inhibition of N1 neuraminidase activity by mouse immune sera, BAL or control antibody Z2B3. Sera from the four mice were pooled for each group. BAL could not be collected from mouse b in the CLEARFLU IN+IP group, mouse a in the S-FLU IN+IP group or mouse a in the VGM group. B. Neutralising antibody response to H7 S-FLU by immune mouse sera, BAL or control antibody MEDI8852. The top two plots show neutralisation of strain-matched A/Hong Kong/225/2017 S-FLU by individual mouse sera or BAL. BAL could not be collected from mouse a in the S-FLU IN+IP group or mouse a in the VGM control group. The bottom plot shows neutralisation of unmatched H7 A/Netherlands/219/2003 and A/Taiwan/1/2017 S-FLU by pooled immune sera.

The CLEARFLU version 3 virus generated CD8+ T-cell responses to nucleoprotein in the lungs and spleen comparable to a strain-matched S-FLU, which have previously been shown to protect against heterologous - including heterosubtypic – challenge in mice and ferrets and partially protect pigs (Figure 10).

**Figure 10.**
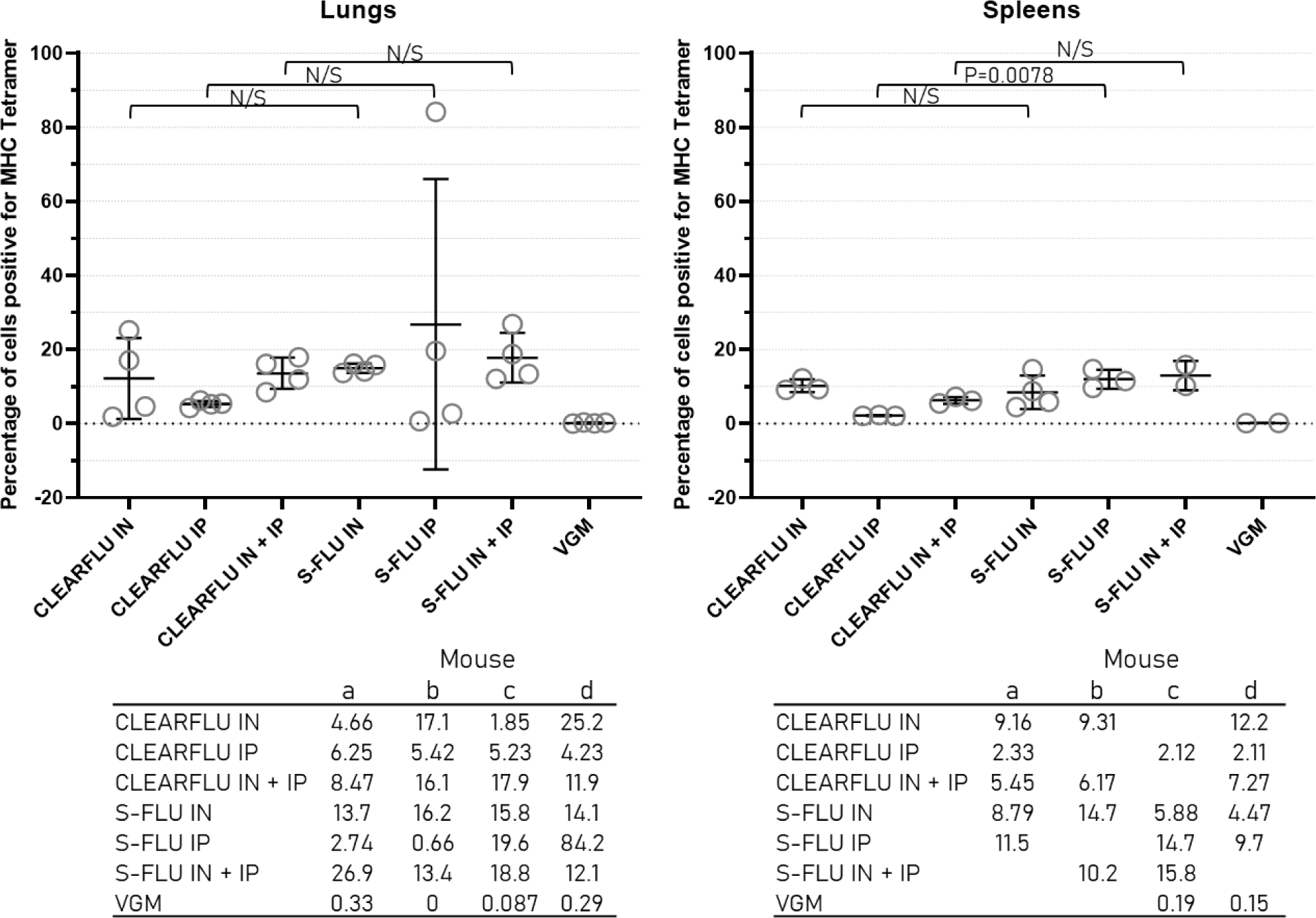
Murine T-cell response to CLEARFLU version 3. Mice were immunised as described in Figure 9. Circles indicate the percentage of CD8+ T-cells positive for MHC tetramer in the lungs and spleen of each mouse a-d in each group. Bars show the mean and standard deviation for each group. For spleens, data could not be collected for 1 individual in the CLEARFLU IN, CEARFLU IP, CLEARFLU IN+IP and S-FLU IP groups and for 2 individuals in the S-FLU IN+IP and VGM control groups. Multiple t tests were used to compare CLEARFLU and S-FLU groups for each administration method. The Holm-Sidak method was used to correct for multiple comparisons. All differences were non-significant at the p=0.05 level except for IP groups in the spleen where p=0.0078.

Overall, these results indicate that CLEARFLU and S-FLU viruses have very similar immunogenicities in mice, with the notable exception that the expression of CLEARFLU hemagglutinins on the surface of cells is able to generate a strong neutralising antibody response after intranasal delivery at low dose, and may therefore provide sterile immunity against strain-matched challenge.

## Discussion

We have described a methodology for designing haemagglutinins inactivated with multiple mutations which can be safely expressed as part of single cycle viruses. We have shown that this approach can be applied to both group 1 (H1) and group 2 (H3 & H7) hemagglutinins without altering expression or antigenicity. In doing so, we have developed a promising vaccine platform for stimulating a combination of broad local T-cell responses and strain-specific neutralising antibodies after intranasal administration. The alternative way to induce neutralising antibody with S-FLU is to combine intranasal administration with a larger dose in the periphery (13,34). A single administration is clearly preferable.

CLEARFLU hemagglutinins expressed within the S-FLU expression cassette were displayed on the cell surface and were immunogenic in mice. Although we identified that some CLEARFLU expression is lost over time during expansion of viral stocks, both viral titre and the proportion of viral clones still expressing CLEARFLU remained high for five rounds of expansion, allowing large quantities of virus to be produced without the need for recloning. While after expansion some viral clones expressed a truncated CLEARFLU segment, we found no mutations in viral clones with a full-length segment, indicating that the CLEARFLU segment was relatively genetically stable. Truncation of the CLEARFLU segment appeared to be associated with a T148A mutation in the neuraminidase which enabled neuraminidase-dependent sialic acid binding and enhancement of cell entry. CLEARFLU viruses overall appeared to be genetically stable, as no mutations were identified elsewhere in the viral genome.

Hemagglutinin-inactivated single cycle influenza viruses like CLEARFLU could be suitable as pre-pandemic as well as seasonal vaccines as they do not encode a viable hemagglutinin that could theoretically reassort with seasonal viruses. The presence of multiple independent mutations scattered throughout the hemagglutinin renders negligible the probability of reversion to pathogenicity and reduces the already low risk of homologous recombination with wildtype hemagglutinins to yield functional new variants (35). As hemagglutinin is a key contributor to antigenic shift, it is particularly beneficial to prevent this segment from acting as a donor sequence for wildtype viruses. Thus an S-FLU expressing a CLEARFLU version 3 H7 haemagglutinin would be theoretically safer than a LAIV encoding a viable H7 sequence (36,37) that could in principle reassort with a seasonal influenza.

Moreover, as the pseudotyping hemagglutinin is the determinant of cell entry but not the antigen driving neutralising antibody responses after intranasal dosing, pseudotyping cell lines can be selected to avoid pre-existing immunity while maximising sialic acid binding. Meanwhile, the multi-mutated CLEARFLU hemagglutinin can be chosen entirely for its immunogenicity. This could be a key advantage over other approaches like LAIV where pre-existing immunity reduces vaccination efficiency against seasonal influenza.

Unlike for many single cycle viruses which lack all or a large part of a viral gene, the full complement of viral antigens should be expressed in RNA, protein and peptide form in the appropriate cellular location after immunisation with CLEARFLU, maximising immunogenicity. In particular, neutralising antibody responses to both hemagglutinin and neuraminidase are desirable as they act independently (38) to limit or abolish virus shedding and onward transmission (39), reduce the risk of pathology in individuals with weak T-cell responses and restrict the opportunity for wildtype virus evolution. CLEARFLU therefore represents a clear advantage over S-FLU and other intranasal vaccine candidates which do not induce neutralising antibody.

## Methods and Materials

### Experimental models

Madin-Darby Canine Kidney-sialyltransferase1 (MDCK-SIAT1) cells generated by transducing MDCK cells with human 2,6-sialtransferase1 (SIAT1) to express higher levels of sialyl-α2,6-galactose moieties (31) were obtained from the European Collection of Authenticated Cell Cultures (ECACC 05071502). Human Embryonic Kidney (HEK) 293T cells (HEK cells stably transduced to express SV40 Large T antigen) were obtained from the William Dunn School of Pathology, Oxford (Ervin Fodor). All cells were grown in D10 (10% FCS + 2 mM L-glutamine + 100 units/ml Penicillin + 100 μg/ml Streptomycin in DMEM) at 37°c with 5% CO2 and passaged when confluent. MDCK-SIAT1 cells were harvested by incubating with Trypsin-EDTA for 10 minutes and centrifuging in 50ml of D10. 293T cells were harvested using 2mM EDTA in phosphate buffered saline (PBS) and spun in 50ml PBS. Pellets were resuspended in D10 for passaging, viral growth medium (VGM) for viral assays or at about 10 million cells/ml in freezing medium for storage.

C57BL/6 and BALB/c female mice were obtained from Envigo RMS Inc, Bicester and maintained in individually ventilated cages at the Biomedical Services facility at the John Radcliffe hospital, Oxford. They were 6-8 weeks old at the start of experiments. Experiments were conducted under Project License number PBA43A2E4 in accordance with the 3Rs.

### Molecular studies

CLEARFLU hemagglutinins were designed by incorporating inactivating mutations into wild-type hemagglutinin sequences from A/England/195/2009 (H1), A/Hong Kong/5738/2014 (H3) and A/Hong Kong/125/2017 (H7). DNAs were human codon-optimised. Restriction enzyme cloning sites for NotI and EcoRI were added to either end of the sequence to facilitate cloning into vector pHR-SIN or the S-FLU cassette for mammalian or viral expression respectively. All sequences were ordered from GeneArt and resuspended in 50μl 10mM Tris/0.1mM EDTA. Sequences are available in the supplementary material.

For subcloning into pHR-SIN (38) or the S-FLU expression cassette, 2μg of GeneArt plasmid was digested using EcoRI and NotI in NEBuffer 3.1. Digestion products were run on gels containing 0.7% Agarose and 0.5μg/ml Ethidium Bromide in TAE buffer. Samples were loaded using Orange G gel loading dye. Bands were imaged and extracted under UV light and purified using a QIAquick Gel Extraction Kit from Qiagen according to the manufacturer’s protocol. Inserts were ligated into new vectors in a 30μl reaction containing 1.5μl T4 DNA ligase, 3μl T4 DNA ligase buffer, 3μl pre-digested and phosphatase treated vector and 15μl of insert. The reactions were incubated at room temperature for at least 1 hour.

All plasmids were grown in DH5α E.coli. 2μl of plasmid was added to 5μl of DH5α ultracompetent cells in Eppendorf tubes on ice. After a 40-minute incubation, a heat shock was performed for 30 seconds at 42°c, followed by incubation on ice for 1 minute. 1ml of LB was added before incubating at 37°c for 1 hour and then spreading the broth on LB-Agar plates containing ampicillin. After overnight incubation at 37°c, colonies were picked into 13ml LB containing ampicillin for a further overnight incubation at 37°c on a 200rpm shaking platform. 0.9ml of the broth was stored at −80°c in Hogness solution for future use. The remaining cells were then pelleted by centrifugation at 3000rpm for 45 minutes and the supernatant removed. Plasmids were extracted from the pellets using a QIAprep Spin Miniprep Kit. To check pHR-SIN plasmids for correct insert length, 20μl digests with 2μl plasmid DNA and 0.5μl each of NotI and EcoRI were set up. Phusion PCR (Thermo Scientific) was performed as recommended by the manufacturer. PCR products were run on 0.7% agarose gels containing ethidium bromide as described for digestion products.

Agilent QuickChange Lightning Site-Directed Mutagenesis kits were used to introduce specific mutations by PCR into the hemagglutinin sequence of A/Hong Kong/125/2017 to generate versions with a single inactivating mutation set. The sequence was subcloned into vector pcDNA3.1- as described above for mutagenesis, and then subcloned into vector pHR-SIN afterwards using the same restriction sites. The bacteria provided were grown up in the same conditions as described for DH5α E.coli.

### Transfection and transduction of cell lines

Multi-plasmid transfection of HEK 293T cells for generating recombinant viruses was performed using Lipofectamine 2000 as described previously (13,9).

MDCK-SIAT1 cells were transduced to stably express hemagglutinin by lentivirus transduction. First, HEK 293T cells were transfected with 1.33μg each of plasmids containing VSV-G, Gag/Pol and the gene of interest in a pHR-SIN vector to produce lentivirus (40). After 48 hours, the supernatant was incubated with MDCK-SIAT1 cells in the presence of 8μg/ml polybrene in D10. After 24 hours, the lentivirus infection was repeated and cells were harvested once confluent.

### Staining cells for fluorescence-activated cell sorting (FACS)

Confluent T175 flasks of MDCK-SIAT1 cells were harvested and spun in D10 at 1400rpm. The cell pellet was resuspended in 1ml of 20μg/ml primary antibody in D10 with 10mM HEPES and incubated on ice for 1 hour. Cells were then washed with 50ml of D10 and resuspended in 1ml of 20μg/ml secondary antibody and incubated on ice for 1 hour. Cells were again washed with D10, spun and resuspended in 7ml of D10 before being passed through a 0.22μm syringe filter and sorted using a BD FACSAria III. Sorted cells were grown up and passaged in T175 flasks to produce cell stocks for subsequent experiments.

### Staining cells for analysis by flow cytometry

MDCK-SIAT1 cells were aliquoted into FACS tubes at approximately 1 million cells per tube. The tubes were spun at 1400rpm and the cells were resuspended in 50μl of 20μg/ml primary antibody diluted in cold FACS wash (1% FCS + 0.01% Azide in PBS). After 1 hour at 4°c, 2ml of FACS wash was added and the cells were spun and resuspended in 50μl of 20μg/ml secondary antibody. After a further hour at 4°c, the cells were washed in FACS wash and resuspended in 300μl of FACS fixative (1% FCS + 1% Formalin in PBS). Cells were analysed using a CyAn ADP flow cytometer by Beckman or an Attune NxT flow cytometer by Invitrogen. Data was analysed using FlowJo v10.6.1. Forward and side scatter were used to gate for single cells only.

### Staining cells for plated viral assays

Confluent MDCK-SIAT1 cells in 96 well plates were stained with 40μl of 1-5μg/ml primary antibody per well for 1 hour at 4°c, washed thoroughly with PBS and then stained with 40μl of 5μg/ml secondary antibody in the same conditions. For staining nucleoprotein, cells were first fixed by incubation with 100μl of 10% formalin for 30 minutes at 4°c and then incubated with 100μl of permeabilization buffer (0.5% Triton X-100) at room temperate for 20 minutes. For stains not involving nucleoprotein, cells were fixed after staining and were not permeabilised. Double stains were performed by incubating with both primary antibodies - one mouse and one human - simultaneously and then both secondary antibodies, as they do not cross-react significantly. Antibodies were diluted in FACS wash. Fixed cells were stored in PBS. Images were taken with a Zeiss fluorescence microscope using 10x magnification. Images were managed using Fiji/Image J. Fluorescence was quantified using a CLARIOstar microplate reader (BMG Labtech).

### Virus titrations

Isolation of viral clones by limiting dilution and quantification by 50% tissue culture infectious dose (TCID50) was performed as described previously (13). Briefly, virus was added to 3E4 MDCK-SIAT1 cells in serial half-log dilutions in a 96-well plate in 200μl volume of VGM with TPCK-trypsin, with 8 replicates for each dilution. After 48 hours, cells were fixed, permeabilised and stained as described above and the dilution at which 50% of wells were infected was calculated using the Reed Muench method (41).

The cell infectious dose (CID50) was determined by adding 3E4 MDCK-SIAT1 cells to serial two-fold dilutions of virus in duplicate rows in a 96-well plate without the addition of trypsin. After overnight incubation, the cells were fixed, permeabilised and stained and the dilution of virus at which 50% of cells in a well were infected was determined by linear interpolation.

## In vivo work

### Immunisation and sample collection

For intranasal immunisation, mice were anaesthetised using 4.5% isofluorane and drops of virus were pipetted onto the nose to be inhaled one by one to a total volume of 50μl. For intraperitoneal immunisation, 500μl of virus was injected into the intraperitoneal space using a 0.5ml insulin syringe and a 29G needle. Mice were humanely killed by exposure to slowly rising concentration of CO2 and death was confirmed by cervical dislocation. Bronchoalveolar lavage (BAL) was obtained by nicking the trachea and using a syringe to wash 1ml of sterile PBS through the lungs three times. Blood was harvested by cardiac puncture using a 23-25G needle and a 1 or 2ml syringe and stored in a BD microtainer with SST gel for 30 minutes to allow clotting before spinning at 10000rpm for 5 minutes. Serum was collected from above the interface gel and heat-inactivated at 56°c for 30 minutes. Spleens were cut into small pieces and passed through a 70μm cell strainer mesh to make single cell suspensions in R10 (10% FCS + 2mM L-glutamine + 100 units/ml Penicillin + 100μg/ml Streptomycin in RPMI-1640). Lungs were minced and treated with 4ml of enzyme solution (2mg/ml Collagenase IV + 200 units in RPMI-1640) for 30 minutes at 37° before being passed through a 70μm cell strainer mesh. The suspension was spun at 1200rpm for 5 minutes and the cell pellets were treated with RBC lysis buffer (Qiagen) for 10 minutes and then washed with R10. The cell pellets were resuspended in 5ml of 40% Percoll in a 15ml tube. A 5ml layer of 80% Percoll was laid down at the bottom of the tubes using a Pasteur pipette before spinning the cells at 2000rpm for 25 minutes using a slow deceleration rate. Leukocytes were collected from the Percoll interface and washed in R10 twice before being resuspended in R10.

### Microneutralisation assays

were performed as described previously (13) with slight modifications. Sera were diluted 1 in 20 before use, BAL samples were not prediluted. For some experiments, sera from animals in the same group were pooled. S-FLU viruses with known neutralisation profiles were used as sources of hemagglutinin and were titrated before use to ensure maximum signal. Briefly, sera or BAL were set up in serial two-fold dilutions in 50μl PBS in a 96-well plate and incubated with 50μl S-FLU virus for 2 hours at 37°c before adding 3E4 MDCK-SIAT cells in 100μl VGM. After overnight incubation at 37°c, eGFP fluorescence was read.

### Enzyme-linked lectin assays (ELLA)

were performed to assay inhibition of neuraminidase activity by immune sera or BAL (42, 43; modified by 15). S-FLU viruses were used as sources of active neuraminidase and were selected to avoid interference by neutralising antibodies to hemagglutinin. 96-well ELISA plates were coated in fetuin by incubating in 50μl of fetuin solution overnight at 4°c. Doubling dilutions of sera or BAL were set up in 60μl, to which 60μl of S-FLU virus was added for incubation at 37°c for 2 hours. Fetuin plates were washed four times with PBS before adding 100μl of S-FLU and serum/BAL mix to each well and incubating for 18 hours at 37°c. After washing four times with PBS, 50μl PNA-HRP (peanut agglutinin conjugated to horseradish peroxidase) solution was added for a 1.5 hour incubation. After washing four times with PBS, 50μl of peroxidase substrate solution (OPD) was added and the plates were developed for approximately 15 minutes before stopping the reaction with 50μl of 1M H2SO4. Absorbance at 492 nm was read immediately using a CLARIOstar microplate reader.

### Detection of NP specific T cells

Approximately 1 million homogenised splenocytes or lung cells were added to each well of a 96 well plate and washed with R10. MHC class I tetramers (NP_366-374_ ASNENMETM labelled with APC) were diluted 1 in 200 in R10 and 200μl was added to each well. After 30 minutes at 37°c, the cells were washed with PBS. 25μl of Zombie Violet was used at 1:1000 in PBS to stain dead cells by incubating in the dark at room temperature for 10 minutes. 25μl of an antibody mix was added to each well to simultaneously stain for surface markers CD8 (APC-Cy7 53-6.7), CD44 (Alexa Fluor 488 IM7), Cd11b (Brilliant Violet 421 M1/70) and B220 (Brilliant Violet 421 RA3-6B2), with each antibody diluted 1 in 200 in FACS buffer. After 30 minutes on ice in the dark, the cells were washed with PBS and fixed using 100μl of cold 5% formalin for 10 minutes at 4°c, and then washed again and resuspended in PBS. CD8+ T-cells positive for MHC tetramer were analysed using an Attune NxT flow cytometer (Thermo Fisher Scientific) and FlowJo v10.6.1. Ultracompensation beads were used to account for overlapping emission spectra.

### Data availability

Data generated in this study are available from the corresponding author on request. The products generated can be made available to other laboratories for research use.

## Acknowledgements

This research was funded by the Biotechnology and Biosciences Research Council Doctoral Training Partnership (BBSRC DTP) at the University of Oxford (Grant number: BB/M011224/1)

